# Validation of a Microfluidic Device Prototype for Cancer Detection and Identification: Circulating Tumor Cells Classification Based on Cell Trajectory Analysis Leveraging Cell-Based Modeling and Machine Learning

**DOI:** 10.1101/2024.08.19.608572

**Authors:** Rifat Rejuan, Eugenio Aulisa, Wei Li, Travis Thompson, Sanjoy Kumar, Suncica Canic, Yifan Wang

**Affiliations:** Department of Mathematics and Statistics, Texas Tech University, Lubbock, TX, USA; Department of Chemical Engineering, Texas Tech University, Lubbock, TX, USA; Department of Mathematics, University of California Berkeley, Berkeley, CA, USA

## Abstract

Microfluidic devices (MDs) present a novel method for detecting *circulating tumor cells* (CTCs), enhancing the process through targeted techniques and visual inspection. However, current approaches often yield heterogeneous CTC populations, necessitating additional processing for comprehensive analysis and phenotype identification. These procedures are often expensive, time-consuming, and need to be performed by skilled technicians. In this study, we investigate the potential of a cost-effective and efficient hyperuniform micropost MD approach for CTC classification. Our approach combines mathematical modeling of fluid-structure interactions in a simulated microfluidic channel with machine learning techniques. Specifically, we developed a cell-based modeling framework to assess CTC dynamics in erythrocyte-laden plasma flow, generating a large dataset of CTC trajectories that account for two distinct CTC phenotypes. Convolutional Neural Network (CNN) and Recurrent Neural Network (RNN) were then employed to analyze the dataset and classify these phenotypes. The results demonstrate the potential effectiveness of the hyperuniform micropost MD design and analysis approach in distinguishing between different CTC phenotypes based on cell trajectory, offering a promising avenue for early cancer detection.

**Author summary:** Early detection is currently the most effective method to combat cancer, as it maximizes treatment options and improves potential survival rates. However, the cost of early detection presents a significant barrier, limiting access for underrepresented groups and discouraging industrial partners from investing in the research and development of screening devices. This study provides an in-silico conceptual validation for the development of an innovative hyperuniform microchip designed to identify circulating tumor cells (CTCs) without the need for biomarker labeling. We created a cell-based modeling framework to examine CTC dynamics in erythrocyte-laden plasma flow, producing an extensive dataset of CTC trajectories that reflect two distinct CTC phenotypes. Two machine learning architectures were utilized to analyze this dataset and classify the phenotypes. The results demonstrate the potential effectiveness of the hyperuniform micropost MD design and analysis approach in distinguishing between different CTC phenotypes based on cell trajectory, offering a promising and cost-effective method for early cancer detection.

## Introduction

The American Cancer Society projects that by 2050, the number of people with cancer will increase by 77%, with 1 in 5 individuals developing some sort of cancer in their lifetime [1]. Metastasis is a technical term that refers to the spreading of cancer throughout the body; some cancers metastasize late, while others spread years before the primary tumor is detected [2, 3]. Cancer is considered metastasized when tumor cells migrate from the primary site to secondary locations, forming new tumors in potentially distant parts of the body.

Tumor cells can migrate in several ways; those that migrate via the blood, or lymphatic, vessels are called *circulating tumor cells* (CTCs). The detection of CTCs can help clinicians determine the stage and severity of cancer progression and aid in the tailoring of a patient-specific plan of treatment [3–5].

Among the various options for CTC detection [6–9], the use of microfluidic devices (MD) confers several advantages. MDs can enhance CTC detection through the use of targeting methods; examples include 3D channels coated with specific antibodies or incorporating close-proximity magnetic or electric fields, that increase the probability of adherence for specific CTC phenotypes. MDs are often constructed from transparent materials that can ease the task of characterizing labeled CTC using visual, or image based, inspection. Finally, the form factor of a MD is typically compact and manufacturing is cost-effective; these features make MDs a natural choice for clinical deployment.

Designing a MD capable of isolating a specific, homogeneous population of CTCs is an important research problem. More specifically, CTCs can exhibit substantially different surface markers, genertic mutations and molecular profiles [10–13]. Variations in CTC characteristics can alter their propensity to initiate the formation of secondary tumors [14]. The general isolation of CTCs, from background objects such as plasma, leukocytes and erythrocytes, is an active area of research [6–9, 15, 16]. However, a limitation of several contemporary approaches is that, while they do isolate CTCs, the isolated CTC population is still heterogeneous [17]. Identifying important subpopulations from an isolated sample often necessitates additional processing, such as applying an immunofluorescent stain or performing downstream genomic or transcriptomic analysis [10, 12], both of which can be time-consuming or costly.

A special class of hyperuniform (HU) micropost obstacle geometry microfluidic devices (HUMDs) have been investigated for their efficacy in the label-free, cost-effective identification of CTCs [13, 18], as illustrated in Figure 1. A HU configuration is a specialized spatial arrangement in which local micropost characteristics are heterogeneous, or random, whereas the global structure is regular, or homogeneous. The fundamental hypothesis of a HUMD is that small-scale structural heterogeneity will generate localized flow patterns and distinct trajectories for CTCs based on properties related to their metastatic profile. For example, localized differences in flow patterns, varying CTC size/shape and elasticity, a property closely related to CTC surface marker density should result in distinct trajectories within a HUMD. Taken together, the HUMD hypothesis is that by analyzing the CTCs’ trajectories through a HUMD, we may gain invaluable information about CTCs metastatic propensity without the need for downstream omics analyses or immunofluorescent labeling.

**Fig 1.**
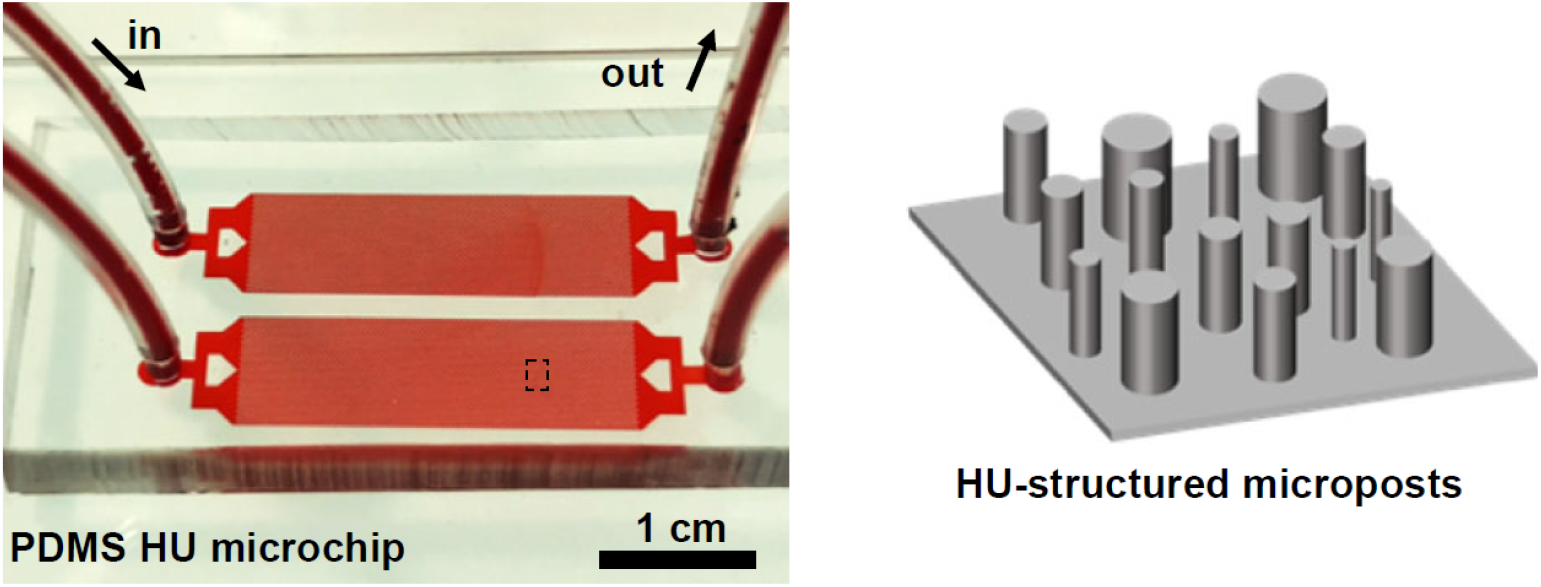
The Hyperuniform microfluidic device includes a schematic illustration depicting the micro-post structure.

This manuscript explores the hypothesis of using a HUMD for CTC classification by integrating mathematical modeling of fluid-structure (cell) interactions in a simulated HUMD with machine learning techniques. This study has two objectives. The first objective is to assess the characteristics of CTC dynamics in erythrocyte laden plasma flow. More specifically, we will examine the interplay between plasma flow, erythrocyte shear stress, and erythrocyte morphology, and how these factors influence CTC dynamics. We will conduct a computational analysis of the role of shear thinning on CTC dynamics through HU microchannels, and investigate how CTCs’ material properties affect their trajectory through an HUMD. The second objective is to generate a CTC trajectory dataset considering two different phenotypes of CTCs based on in-silico simulation, and then apply machine learning techniques to analyze the data and perform classification. This second objective serves as an in-silico validation of our hypothesis for designing a novel early cancer detection microfluidic chip. The manuscript is structured as follows: In the Models section, we describe our mathematical modeling approach. The Results section begins with benchmark validations of our model, showing that it accurately captures the physics of erythrocyte-laden plasma flow. We then present a series of in-silico simulation results that replicate observations from wet-lab experiments. Finally, we show two distinct machine learning architectures, based on CNN and RNN, achieved an accuracy of 84% in classifying CTC phenotypes based on cell trajectory data. This demonstrates the potential effectiveness of our hypothesis regarding the HUDM design and analysis approach for distinguishing between different CTC phenotypes.

## Results

In this section, we conducted several benchmark tests on the Fluid Structure Interaction (FSI) model (as stated in the **Models** section), ranging from simulating RBC motion under various flow conditions to 3D channel flow to study the shear thinning effect of blood. The code implementation is based on the open-source C++ library [19].

### In-silico replication of the known RBC dynamics in a 3D narrow channel flow

We conduct a full 3*D* benchmark test building upon a previous 2*D* study on simulating RBCs suspended in plasma flowing through a rigid cylindrical channel [20]. The channel has a length of 90 *µm* and a radius of 4.5 *µm*. Each RBC has a diameter of 8 *µm*. Initially, the RBCs are evenly spaced at a distance *d*_*RBC*_ from each other, oriented perpendicular to the flow direction, as illustrated in Figure 2. Various hematocrit (Hct) levels, ranging from 0.1 to 0.49, were considered by adjusting *d*_*RBC*_ from 22 *µm* to 4.5 *µm*. For instance, at Hct = 0.1, *d*_*RBC*_ is set to be 22 *µm*. The plasma viscosity is set to 1.0 × 10^*−*3^ Pa *·* s and the density to 1.0 × 10^3^ kg/m^3^. The flow is driven by an external force, resulting in a Reynolds number of 0.1. No-slip boundary condition is assumed between the plasma and the RBCs, as well as between the plasma and the inner cylinder wall. One lattice unit is defined to correspond to a physical length of 0.25 *µm*, resulting in a simulation size of around 0.49 million lattice cells. All other constants, as reported in the **Models** section, are chosen as follows: *k*_*l*_ = 15*K*_*B*_*T, k*_*b*_ = 80*K*_*B*_*T, k*_*v*_ = 20*K*_*B*_*T* and *k*_*a*_ = 5*K*_*B*_*T*.

**Fig 2.**
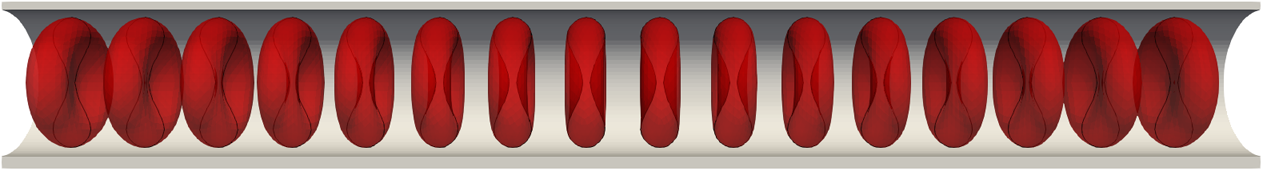
3*D* simulation of blood flow in a long narrow cylindrical channel. 3*D* biconcave RBCs were positioned at varying distances *d*_*RBC*_ to adjust Hct values from 0.1 to 0.4. In this illustrated picture, the Hct=0.4.

Figure 3 illustrates the deformation of red blood cells (RBCs) over time within a flow channel with a Hct level of 0.49. The simulation time *t* is normalized by a characteristic time scale *T*_0_ = *L/u*_0_, where *L* is the characteristic length chosen to be the channel length and *u*_0_ is the characteristic velocity corresponding to the average inlet velocity. To continuously observe the deformation and behavior of the RBCs under steady flow conditions, a periodic flow boundary is used, meaning that when particles exit the outlet of the flow channel, they are reintroduced at the inlet. From the simulation result, we observe that the RBCs initially exhibit a biconcave disk shape, which is their natural form. As time progresses, they gradually deform into a parachute shape. This deformation process is depicted in Figure 3, aligns well with the experimental observations reported in [21]. Beyond the normalized time *t/T*_0_ = 0.60, the flow reaches a steady state, and the RBCs attain their maximum deformation. This steady-state deformation also indicates that the cells have adapted to the flow conditions within the channel. Figure 4 presents the mechanical behavior of a single RBC in blood flow across various Hct levels ranging from 0.1 to 0.49. For better visualization, snapshots of a single RBC at different time points are superimposed within a single channel. This parametric study on Hct revealed that RBCs experienced less deformation at higher Hct level. The degree of deformation was quantified using a relative deformation index 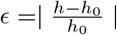, where *h*_0_ is the initial diameter of the RBC (8*µm*) and *h* is the cross-section length of the RBC at time *t*. Relative deformation index *ϵ* remains a constant after the blood flow reaches a steady state. Figure 5 plots the relative deformation index *ϵ* at the steady state (*t/T*_0_ = 1) as a function of Hct level. The results show consistence agreement with the work by Kenichi et al. [20], noting that while Kenichi’s study focuses on a 2D model, our findings are based on a 3D study.

**Fig 3.**
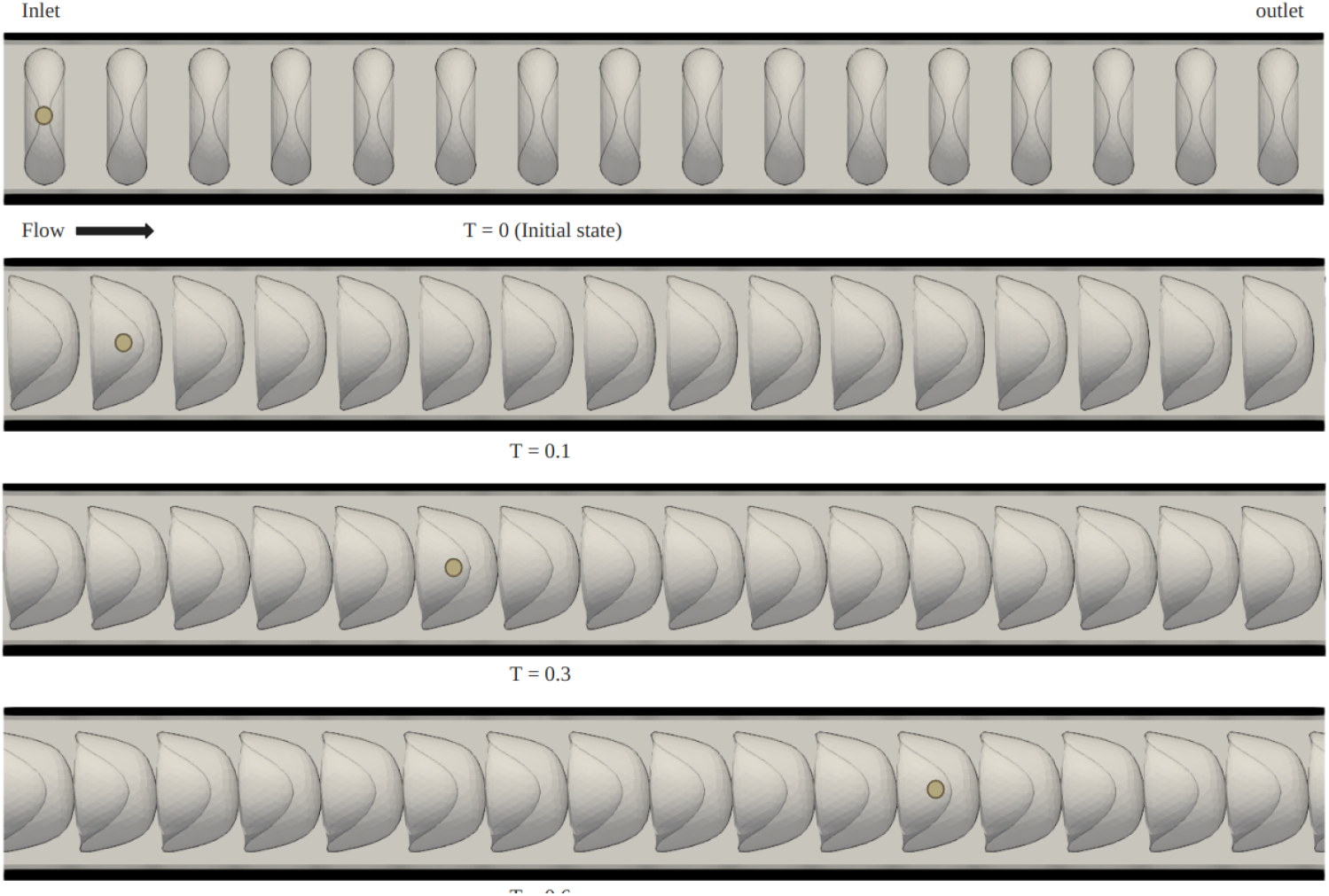
Snapshots showing the RBC deformation in blood flow within a cylindrical channel at different time moments *T* = *t/T*_0_. The same RBC is labeled with a yellow circle for reference. Here *Hct* = 0.49.

**Fig 4.**
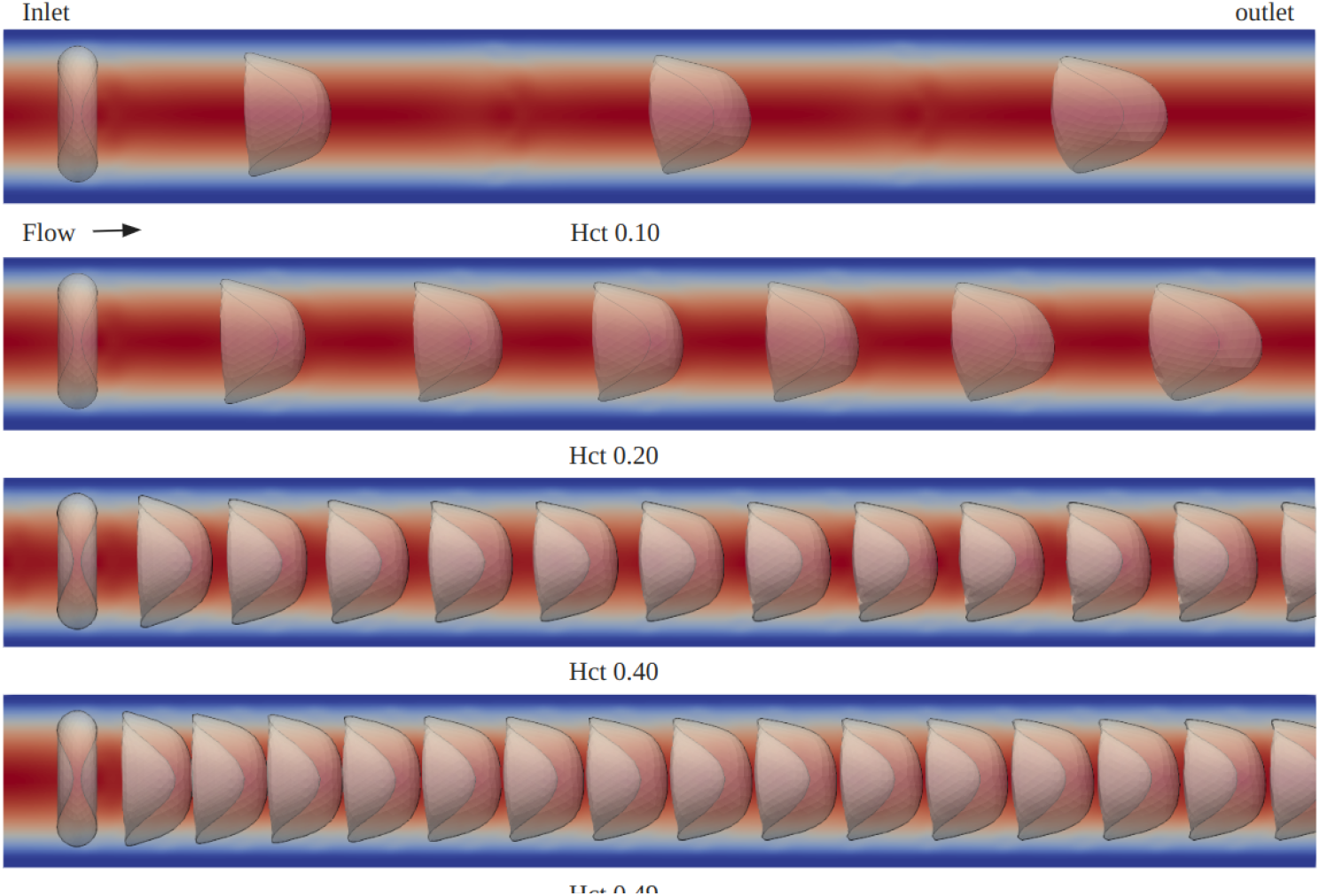
Deformation of a single RBC at various Hct values from 0.1 to 0.49

**Fig 5.**
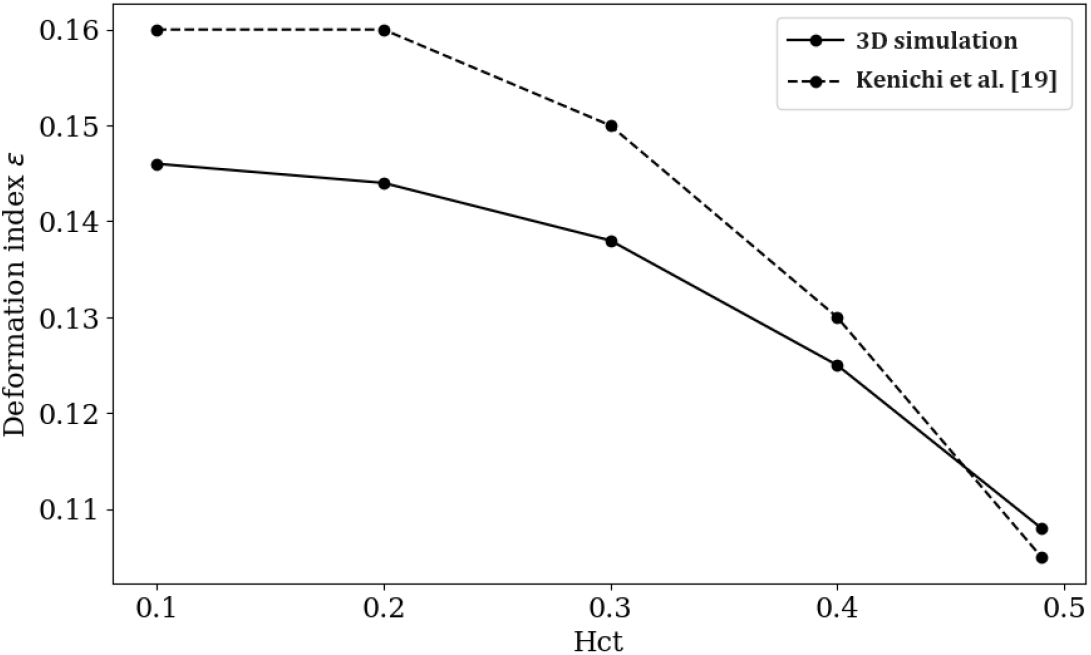
Relative deformation index *ϵ* from time *t/T*_0_ = 1 as a function of Hct. The dashed line represents the results from the 2D model studied by Kenichi et al., while the solid line represents our 3D model.

### In-silico replication of the tumbling, rolling, and tank treading motions of a single RBC in a shear flow

We consider a single RBC positioned at the center of a plasma-filled box region. The fluid region has dimensions of 20 × 20 × 10 *µm*^3^, as illustrated in Figure. 6. The fluid is subjected to a horizontal shearing flow, driven by two parallel plates moving in opposite directions at a velocity of *v*_0_. Given the distance between the top and bottom plates is *d* = 10*µm*, the shear rate is determined by 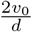. The physical length of the lattice grid is Δ*x* = 0.5 × 10^*−*6^*µm*, and time step is Δ*t* = 0.5 × 10^*−*7^*s*. The diameter of RBC is set to 8 *µm* [22] with all other constant values remaining the same as reported in the **Models** section. To represent different scenarios that might occur in natural blood flow conditions, we explored two different initial orientations of RBC in our study [23–27]. In case (A), we examine the scenario where one of the long axes is aligned parallel to the flow direction, while the other long axis is perpendicular to the flow. In case (B), both long axes are perpendicular to the flow. The initial positions for these two cases are depicted in Figure 6. In the following, we will report the results for cases (A) and (B) separately.

**Fig 6.**
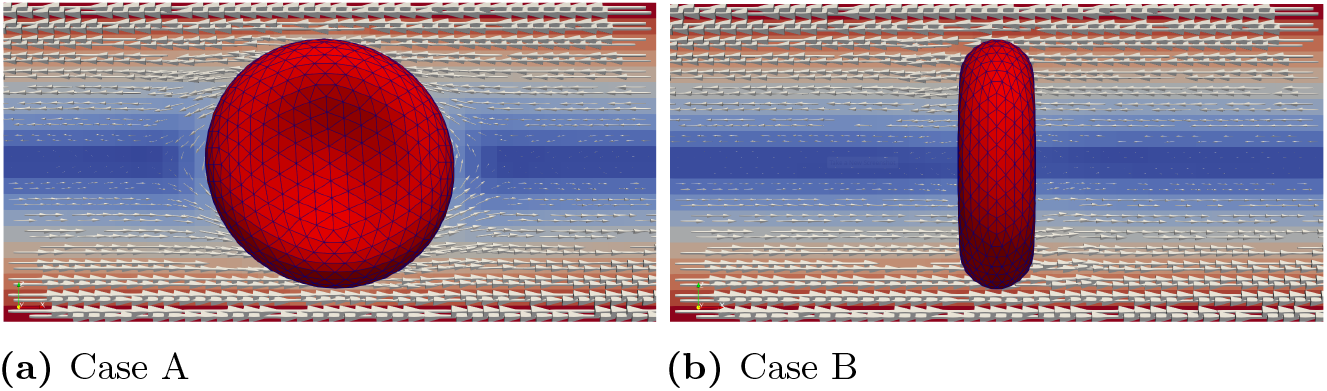
Two different initial RBC orientations regarding the shear flow direction

In case (A), we conducted simulations accoss a range of shear rates, specifically 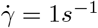, 5*s*^*−*1^, 10*s*^*−*1^, 20*s*^*−*1^, 50*s*^*−*1^, 75*s*^*−*1^, 100*s*^*−*1^. Figure 7 illustrates the scenario with shear rate 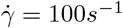. Under this high shear rate flow condition, the RBC exhibits a tank treading motion, and the inclination angle remains fixed. When the shear rate is moderately high (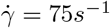 and 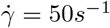), the RBC maintains a tank treading motion with a fixed inclination angle. However, as the shear rate decreases further (below 20^*−*1^), the RBC’s motion gradually transitions to a rolling motion, as shown in Figure 8. (c) and (d). At relatively low shear rates (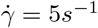 and 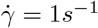), the RBC exhibits a pure rolling motion with minimal or no deformation, as shown in Figure 8. (e) and (f).

**Fig 7.**
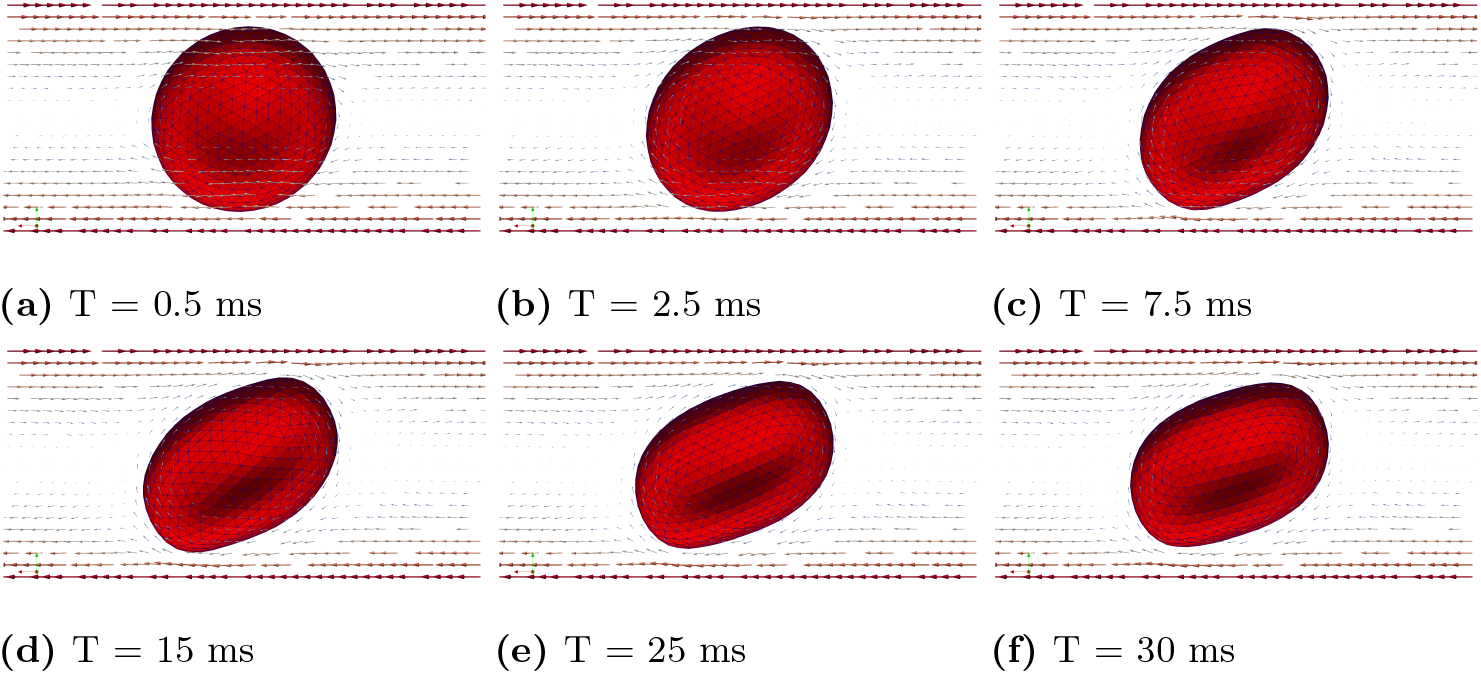
Case (A): Snapshots of velocity field and RBC tank treading motion at different times for 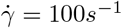. After the initial deformation, the inclination angle remains fixed.

**Fig 8.**
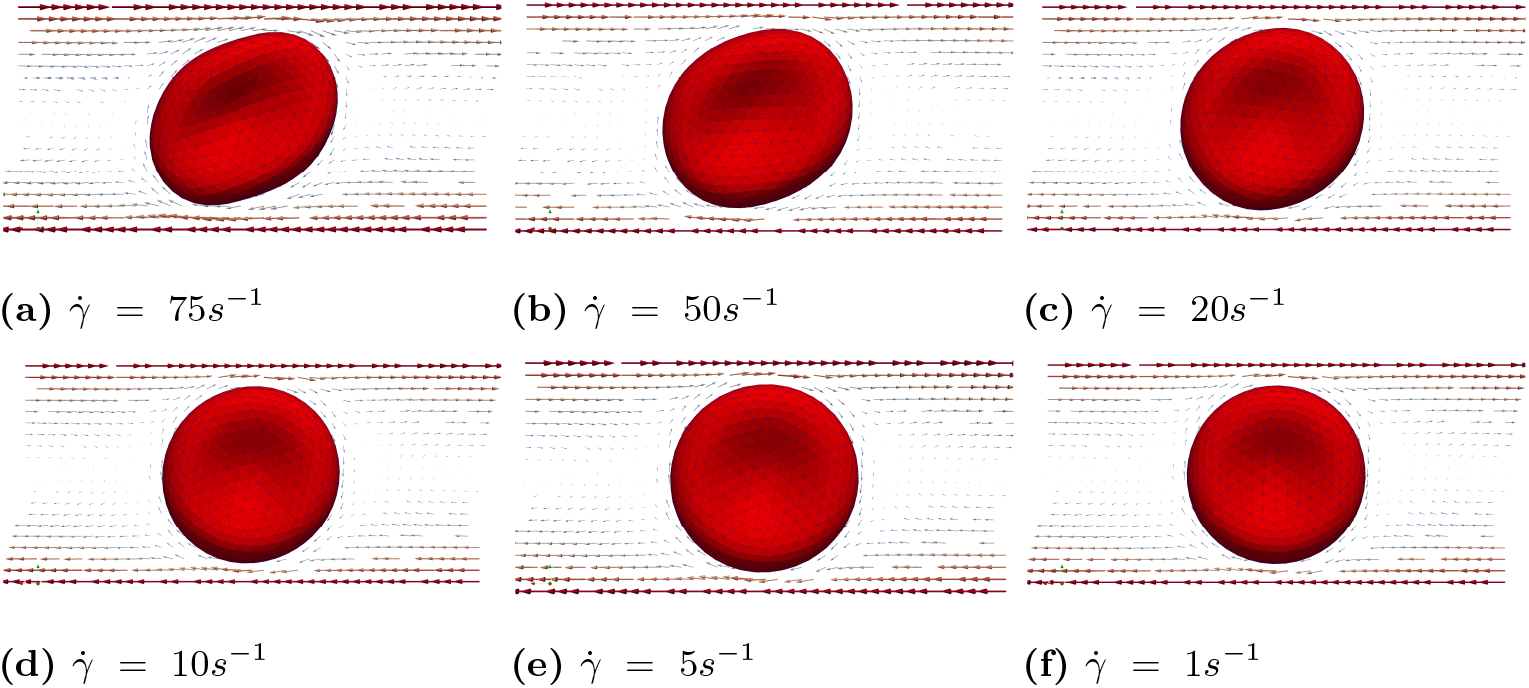
Case (A): Snapshots of velocity field and RBC motion at T = 75*ms* for various shear rates, ranging from 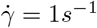, 5*s*^*−*1^, 10*s*^*−*1^, 20*s*^*−*1^, 50*s*^*−*1^, 75*s*^*−*1^.

In case (B), we observed that tank treading motion requires a much higher shear rate. Consequently, we conducted simulations with shear rates 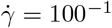, 150*s*^*−*1^, 200*s*^*−*1^, 250*s*^*−*1^, 300*s*^*−*1^, 400*s*^*−*1^, and 500*s*^*−*1^, while keeping all other parameters the same. We identified three distinct types of motion under these shear rates. Under a high shear rates (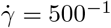 and 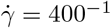), the RBC exhibits a tank treading motion. However, unlike in Case (A), the tank treading motion does not maintain a steady shape due to the inherent rigidity of the biconcave shape and the inertia of the RBC. Figure 9 presents a series of snapshots of the RBC’s cross-sectional view at different times, highlighting the complex dynamics of its motion under high shear rates. When shear rates are between 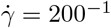 to 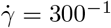, the RBC exhibits a mixed of tank treading and tumbling motions. This is illustrated in Figure 10, which shows snapshots of the RBC’s cross-sectional view at different time points under the shear rate 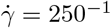. At relatively lower shear rates (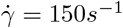 and 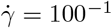), the RBC exhibits only tumbling motion, as shown in Figure 11.

**Fig 9.**
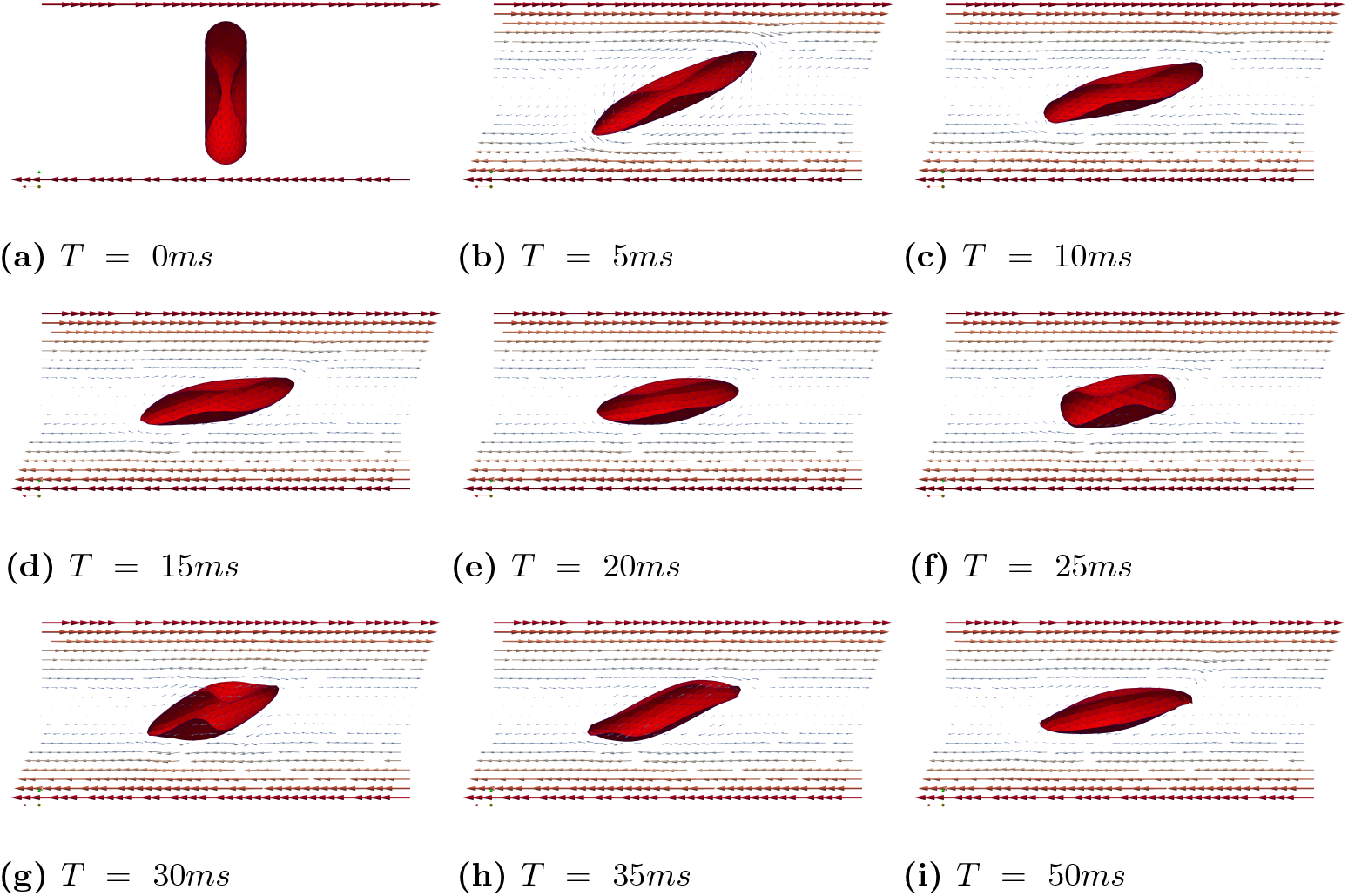
Case (B): Snapshots of the RBC showing tank treading motion at different times under the shear rates 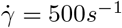.

**Fig 10.**
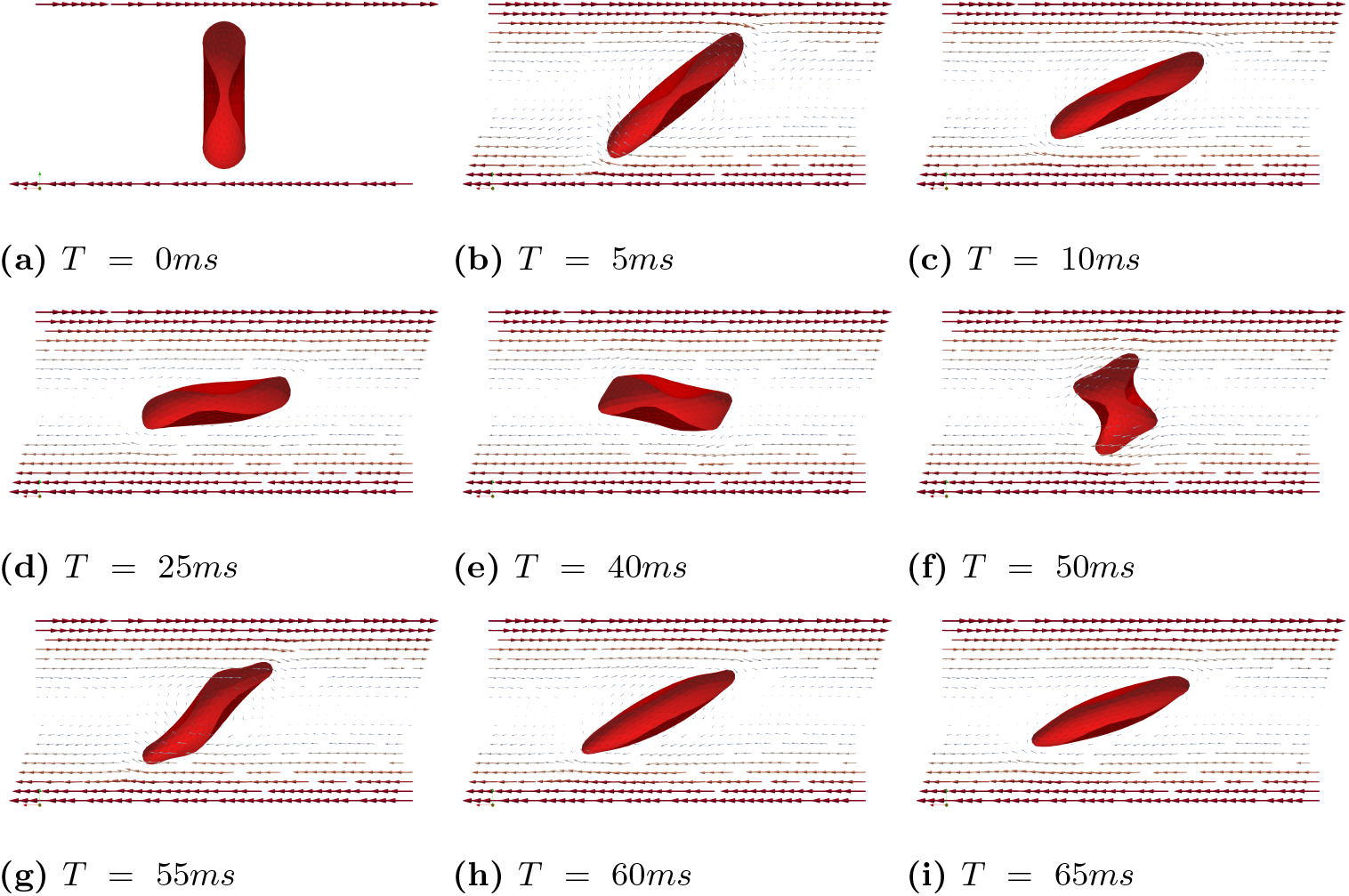
Case (B): Snapshots of the RBC showing a mixed tank treading and tumbling motion at different times for 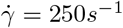.

**Fig 11.**
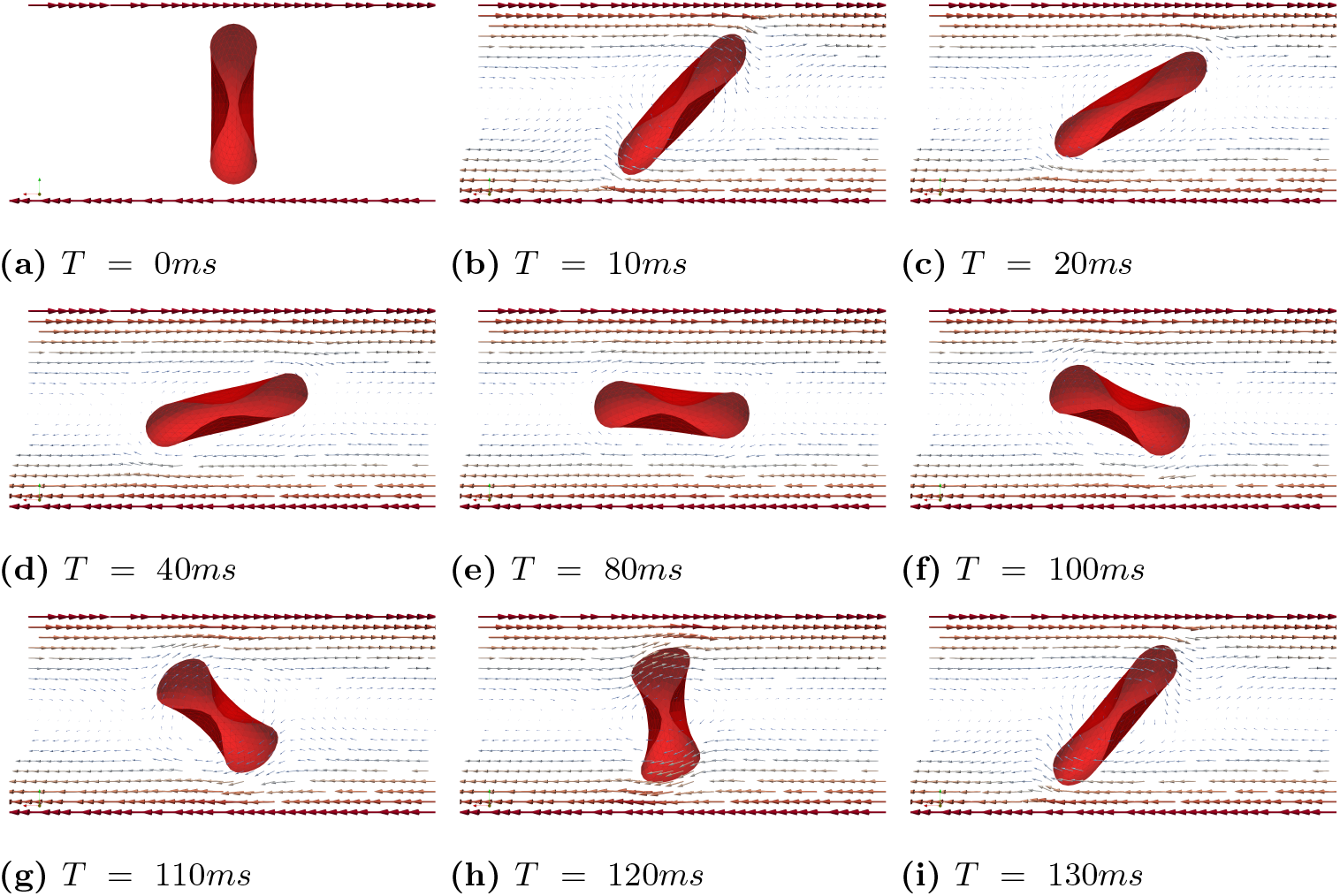
Case (B): Snapshots of the RBC showing tumbling motion at different times for 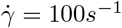.

### In-silico replication of shear thinning effect of blood flow

Blood exhibits non-Newtonian fluid behavior, largely influenced by the rheological properties of RBCs. The apparent viscosity of blood changes under different hemodynamic conditions, such as varying Hct levels and shear rates [28]. For example, at a constant Hct level, the apparent viscosity of blood decreases as the shear rate increases (blood flows more easily when subjected to higher force or stress) [29], a phenomenon known as the shear thinning effect of blood. Under the identical flow conditions (same external force or pressure gradient), a higher Hct level leads to a higher blood viscosity, making the shear thinning effect more pronouced. Conversely, a lower Hct level diminishes the shear-thinning effect until it eventually disappears [30, 31].

To demonstrate that our model effectively captures the shear-thinning phenomena influenced by both hematocrit levels and shear rates, we conducted the following simulations. We simulated blood flow through a channel with dimensions 80 × 80 × 200*µm*^3^, which contains a wedge with dimensions 80 × 40 × 66.67*µm*^3^. We analyzed the relationship between the average shear stress and the shear rate. The lattice grid was set to Δ*x* = 0.5 × 10^*−*6^*µm* and the time step is Δ*t* = 0.5 × 10^*−*7^*s*. All other parameters were kept the same as described in **Models** Section. In the first test, we investigated the blood’s shear-thinning behavior at three different Hct levels: 0 (pure plasma), 0.1, and 0.4. The blood is driven by an external body force, resulting in an inflow Reynolds number of 5. Once the flow is fully established in the whole channel, we calculated the shear stress on the top surface of the wedge using the formula 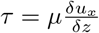, where *u*_*x*_ is the velocity in the longitudinal direction (*x*) and *µ* is the dynamic viscosity. As shown in Figure 12, we present the shear stress for the three cases with the same color scale. It is evident that the case with a lower Hct level (0.1) generates a higher average shear stress, as indicated by the larger yellow regions in Figure 12.(c) than in Figure 12.(b), which corresponds to the higher Hct level (0.4). Furthermore, the case with no red blood cells (*Hct* = 0) results in an even larger yellow area, indicating higher average shear stress on the wedge surface. This test demonstrates that the blood becomes more viscous (resistant to the external force) as the Hct level increases, thereby reducing the shear stress on the top surface of the wedge. To demonstrate the model can effectively capture the shear-thinning effect, we also explored the relationship between the average shear stress on the top surface of the wedge under various shear rates. By maintaining a constant Hct level of 0.1, we varied the inlet flow velocities to achieve different shear rates. Figure 13 illustrates that when the shear rate is below 100 *s*^*−*1^, the relationship between shear rate and shear stress is sub-linear. This shear-thinning behavior is a hallmark of non-Newtonian fluids, commonly observed in blood flow at low shear rates. While at higher shear rates (*>* 100 *s*^*−*1^), the fluid begins to exhibit Newtonian behavior. This aligns with the observations by Ameenuddin et al. [31], who also noted that shear thinning vanishes when the shear rate surpasses 100 *s*^*−*1^.

**Fig 12.**
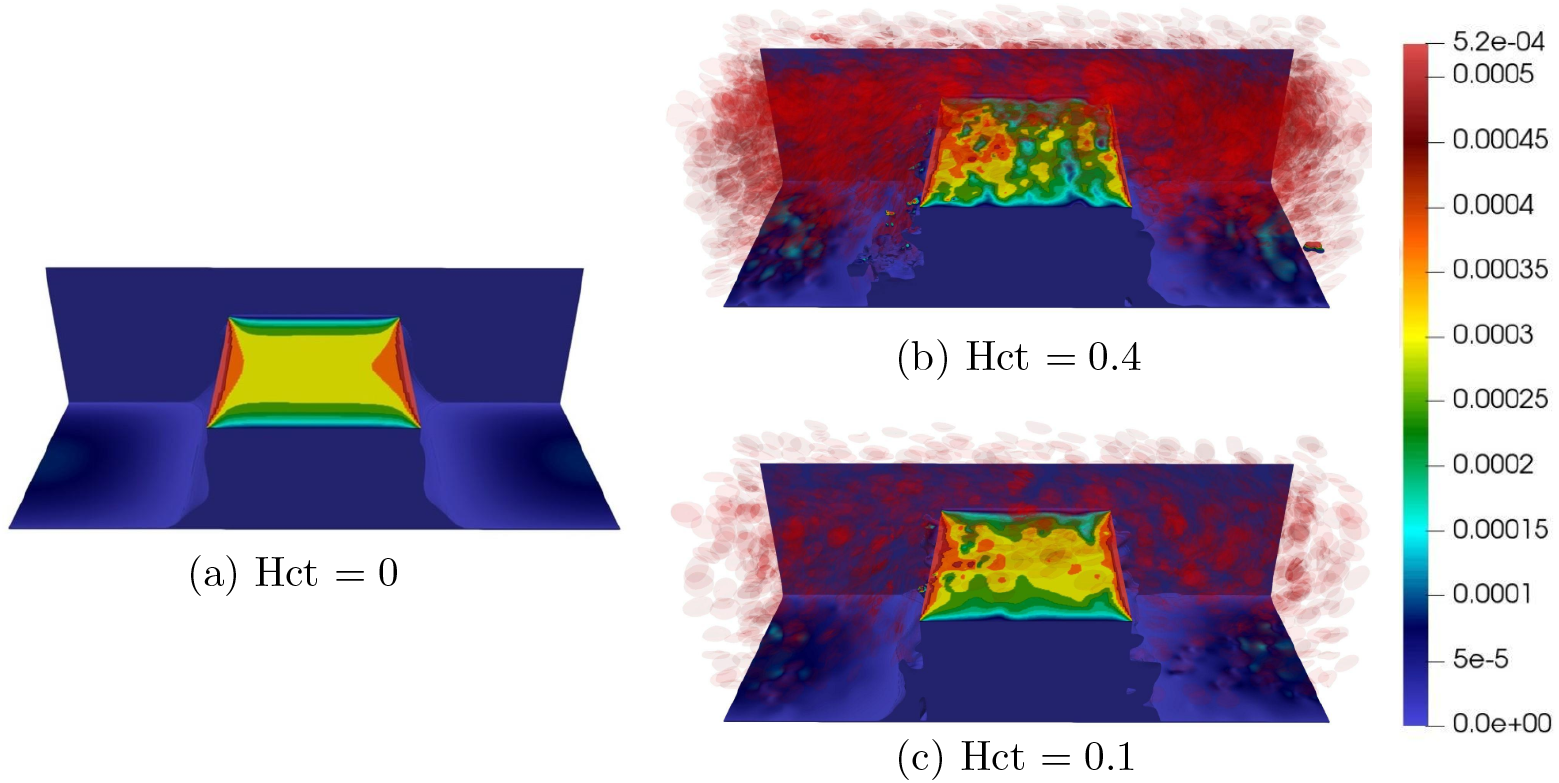
Comparing the shear stress on the top surface of a wedge at different Hct levels: 0 (pure plasma), Hct = 0.1, and Hct = 0.4.

**Fig 13.**
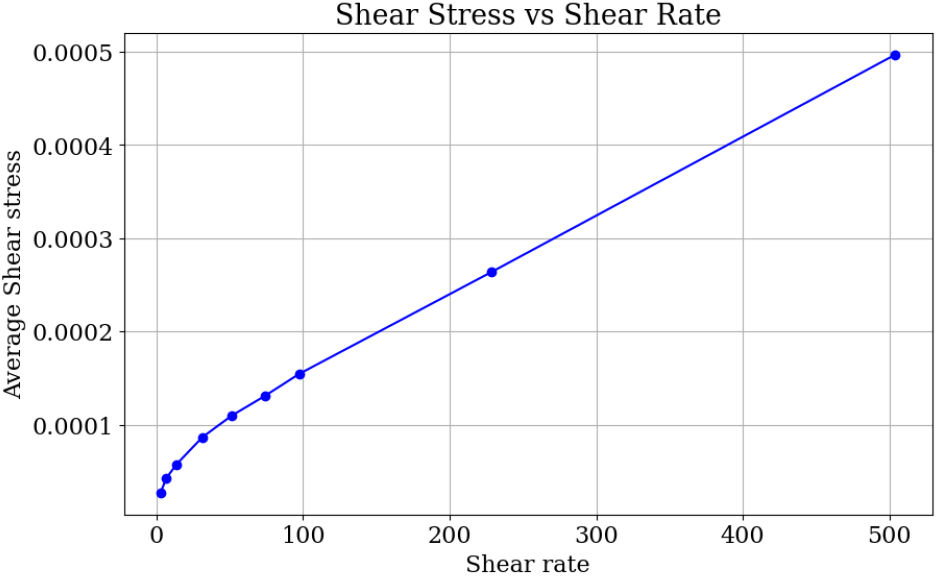
Relation between shear rate and average shear stress (Hct=0.4).

### Modeling CTCs in a micro-fluidic channel and studying the CTC classification

In the following sections, we will use the previously validated cell-based model to investigate the morphology and movement dynamics of circulating tumor cells (CTCs) within the hyperuniform (HU) microfluidic device. The first objective is to use in-silico simulations to replicate the complex dynamics of CTC motion and their interactions within the HU microfluidic device, as observed in experiments. This approach aims to provide deeper insights into CTC behavior in this environment and reduce experimental costs through simulation-based methods. This dual benefit will enhance our understanding of CTC dynamics while making the research process more efficient and cost-effective. The second objective is to generate a set of synthetic CTC trajectory data in silico and investigate the potential of using machine learning algorithms to analyze this data for CTC classification. This second objective is based on the fact that CTCs’ elasticity properties vary across different subphenotypes. These subphenotypes exhibit distinct mechanical characteristics, reflecting their diverse biological origins and states of differentiation. Such variability in elasticity can influence the behavior and trajectory of CTCs within the microfluidic device.

#### Replicating experimental results using numerical simulation

The actual HUMD is designed and manufactured using polydimethylsiloxane (PDMS) through the conventional soft lithography technique in Dr. Li’s lab [13, 32]. The 3D geometry configuration is then utilized for our in-silico simulation. Specifically, we examined a subdomain of the entire channel with dimensions of 300 × 2200 × 25*µm*^3^. The main pipeflow of our modeling process is illustrated in Figure 14. The code is freely accessible, and the link to it is provided in the supporting document.

**Fig 14.**
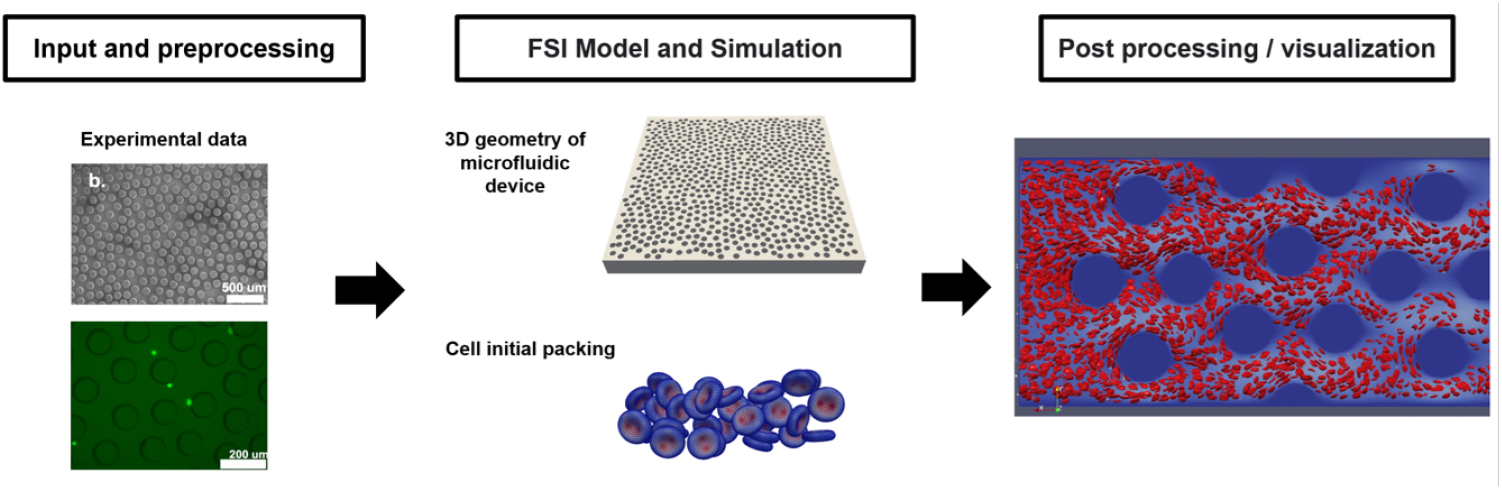
Workflow of in-silico simulation and analysis

#### CTC types and numerical configurations

In the wet lab study, two types of CTCs were examined. In particular, PC3 and LNCap cells harvested from patient samples were fluorescent stained and directed to flow through the HU chip under controlled flow rates. The cells’ trajectories were recorded using high-speed cameras for the subsequent comparison with simulation results. In addition, single-cell RNA sequencing was performed to acquire detailed information about the cell phenotypes and surface EpCAM expression levels. Cell elasticity was measured using atomic force microscopy.

In our in-silico study, similar to our approach with RBCs, the CTC is modeled as an elastic spherical membrane structure. Knowing that CTCs’ morphology and elasticity exhibit dynamic changes among different sub-phenotypes. One of those traits is the EpCAM antigen expression level. For instance, prostate cancer (PC3) cell has an average size of 15±5 *µm* and *∼*50,000 surface EpCAM antigens per cell. In contrast, breast cancer (SKBR3) cell has an average size of 12.5±2.1 *µm* and *∼*600,000 surface EpCAM antigens per cell. The EpCAM expression level can lead to the change of the total elasticity of CTCs up to 10 folds. In the following simulation, we fixed the CTC diameter at 8 *µ* and examined two types of elastic configurations. One cell type was five times stiffer than the other in terms of the elasticity constants *k*_*l*_ and *k*_*b*_, while *k*_*a*_ and *k*_*v*_ remained constant for both types. We tuned the model parameters to accurately replicate the experimental results. Figure 15 demonstrates a good agreement between the trajectories in the experimental image and the numerical simulation.

**Fig 15.**
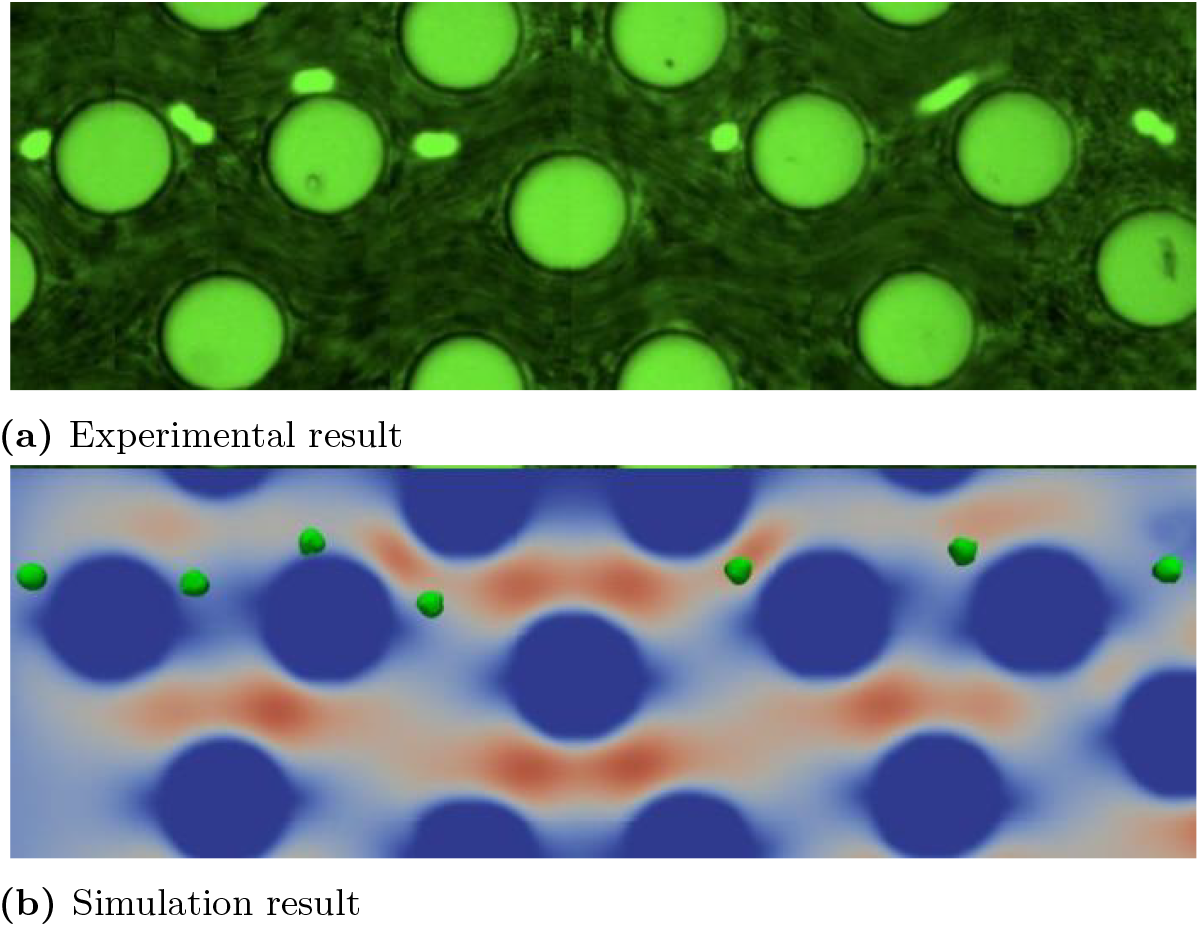
Top: Experimental image showing a PC3 cell traveling through the microfluidic channel, provided by Dr. Li; Bottom: The numerical simulation results based on the identical micro poles configuration. The small green dots represent circulating tumor cells (CTCs), while the large cylindrical shapes represent micro-posts in the HU microfluidic device.

#### Divergence in cell pathways among different cell sub-phenotypes

From the experiments, we observed varying path behaviors among different CTC phenotypes in the HU microfluidic channel, as shown in Figure 16 (a)-(c). We suspect that differences in trajectory might be linked to variations in cell elasticity, which could be leveraged for CTC classification.

**Fig 16.**
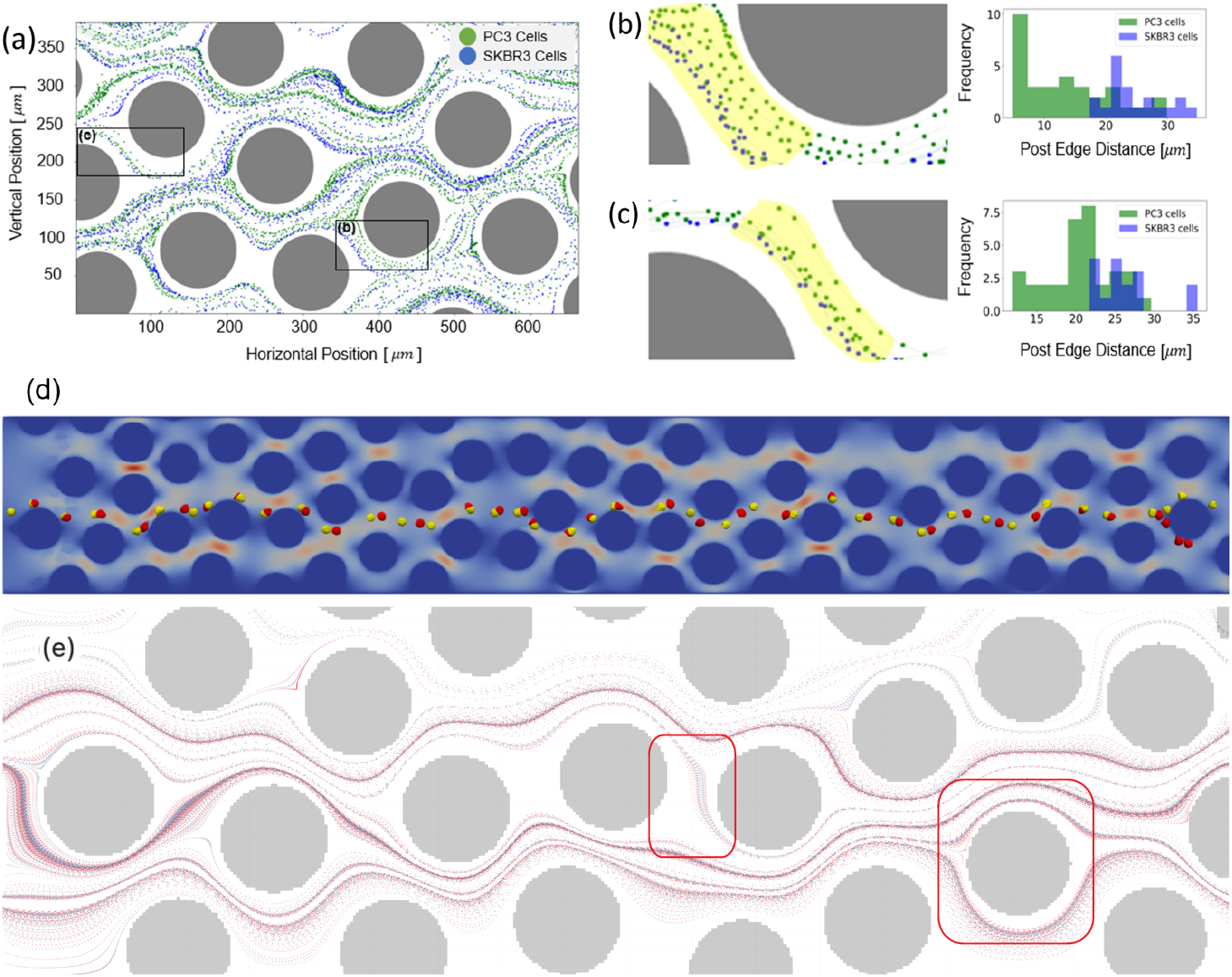
The trajectories of PC3 and SKBR3 cancer cells, obtained from the wet-lab experiment, are displayed in a broader view (a) and two close-up views (b) and (c) highlighted by labeled black boxes. PC3 cells are represented in green, while SKBR3 cells are in blue. We also calculated the distances from each trajectory points to the post edge and plotted the frequencies histograms, courtesy of Li’s lab. The simulation reveals unique paths for two different types of CTCs (red and yellow with distinct elastic properties) starting from the same location initially, as depicted in (d). Cell trajectories of two distinct CTC types (red and blue dots) display clearly different trajectory patterns at those points of interest, as highlighted by the red boxes in (e).

Our preliminary experimental data in Figure 16 showcases the paths of PC3 and SKBR3 within a 350*µm ×* 650*µm* region of the HU microfluidic device. PC3 cells are represented in green, SKBR3 cells in blue, and cylindrical posts in grey. While Figure 16 (a) provides a broader view, Figure 16 (b) and Figure 16 (c) offer a more in-depth local analysis of the two cancer cell lines. The left-hand side of Figure 16 (b) and Figure 16 (c) shows a magnified view from the black boxes marked in Figure 16 (a), while the right-hand side displays the frequency histogram of the post-edge distance of two cancer cell lines. The distance histogram is determined by calculating the trajectory points to the micro-post surface within the yellow highlighted area seen in Figure 16 (b) and Figure 16 (c). Both histogram plots clearly depict a noticeable local distinction in terms of the post-edge distance, leading us to believe that this disparity can be leveraged for CTC classification. This path difference is indicative of varying mechanical and physical properties among cell lines, which may facilitate us to identify the metastatic disparities and phenotypes within different cell lines. Our simulation results confirmed a similar trend, as shown in Figure 16 (d). What even more noteworthy is that the local disparity eventually leads to a global path divergence. As depicted in Figure 16 (d), it’s evident that the paths start to diverge near the end of the channel for two different CTC types. In Figure 16 (e), we highlighted two notable regions where distinct trajectory patterns are observed. For instance, in the left box, red dots are more sparse compared to the denser blue dots. In the box on the right, red dots are positioned closer to the pole surface region than to blue dots. We suspect that the observed differences in trajectory are primarily due to the combined effects of cell deformation and inertial forces. The deformation of the cells as they navigate through the microfluidic channel, along with the inertial forces acting on them, likely contributes to the variation in their movement patterns. This interplay between deformation and inertia may significantly influence how different cell types follow distinct paths.

#### CTC classification based on cell trajectory data

In this section, we present our findings on leveraging machine learning approaches to classify CTC phenotypes. We employed two machine learning models to analyze the cell trajectory data: a Convolutional Neural Network (CNN) and a Recurrent Neural Network (RNN). To generate a sufficient dataset for training these models, we conducted multiple simulations. Specifically, we considered two types of CTCs as described in **CTC types and numerical configurations** section. Initially, the cells were randomly positioned at the inlet, and we recorded their locations at even time intervals of 0.001 second. Over 4000 trajectories were generated, with 2000 corresponding to PC3 cells and 2000 corresponding to SKBR3 cells. For each cell trajectory, the data was stored in six columns: the first three columns correspond to the coordinates of the cell’s center of mass, while the last three columns store the velocity components. The data and the code can be found in the supporting document. We examined this data because these quantities can be easily obtained in an experimental setting using a high-speed camera.

Figure 17 illustrates the architecture of the CNN. Initially, the input data was divided into two components: position and velocity. The position data was processed through a series of convolutional and pooling layers, while the velocity data underwent a separate set of convolutional and pooling layers. The outputs from these two convolutional pathways were then concatenated and fed into a fully connected neural network. Detailed model information can be found in supporting document. Our RNN implementation features multiple Bidirectional layers, each followed by Batch Normalization and Dropout layers. The Dropout layers help prevent overfitting while preserving the output shape. Each Bidirectional layer reduces the sequence dimension progressively. Then, the model concludes with two Dense layers. Detailed information about the model can be found in supporting document.

**Fig 17.**
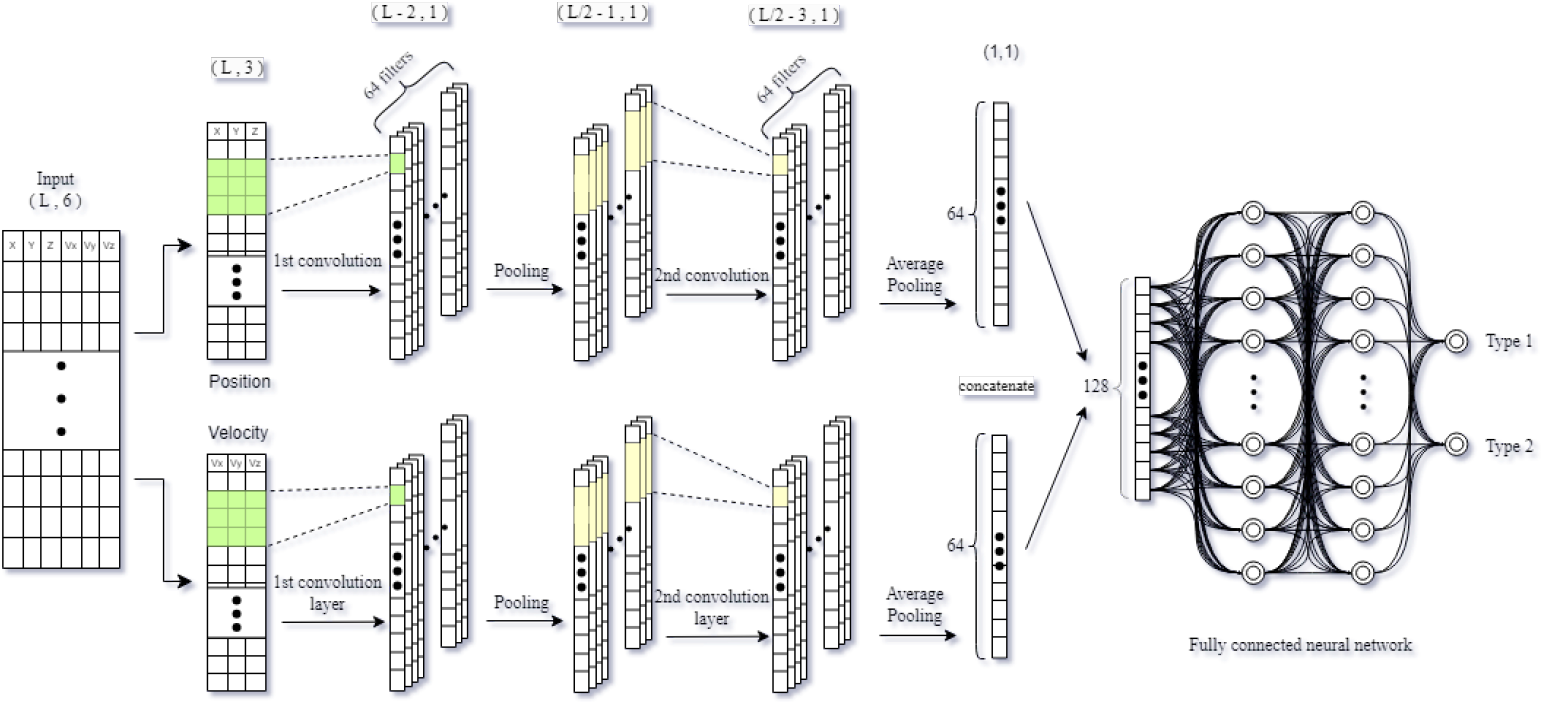
Schematic diagram showing the architecture of the convolutional neural network

Figure 18 shows the training, validation, and test accuracies, as well as the ROC curves for both models. Although the models utilize different techniques for processing input data, their comparable test accuracy and ROC curves suggest that both are likely to deliver similar predictive performance on new data. With both models achieving an accuracy of approximately 84%, our proposed approach to CTC classification demonstrates promising potential. Moreover, this level of accuracy suggests that our method can effectively differentiate between various CTC phenotypes even within a short channel region. Given a full-length HUMD channel, the accuracy is expected to improve further. These results highlight the potential of the proposed HUMD for practical application in cancer diagnostics.

**Fig 18.**
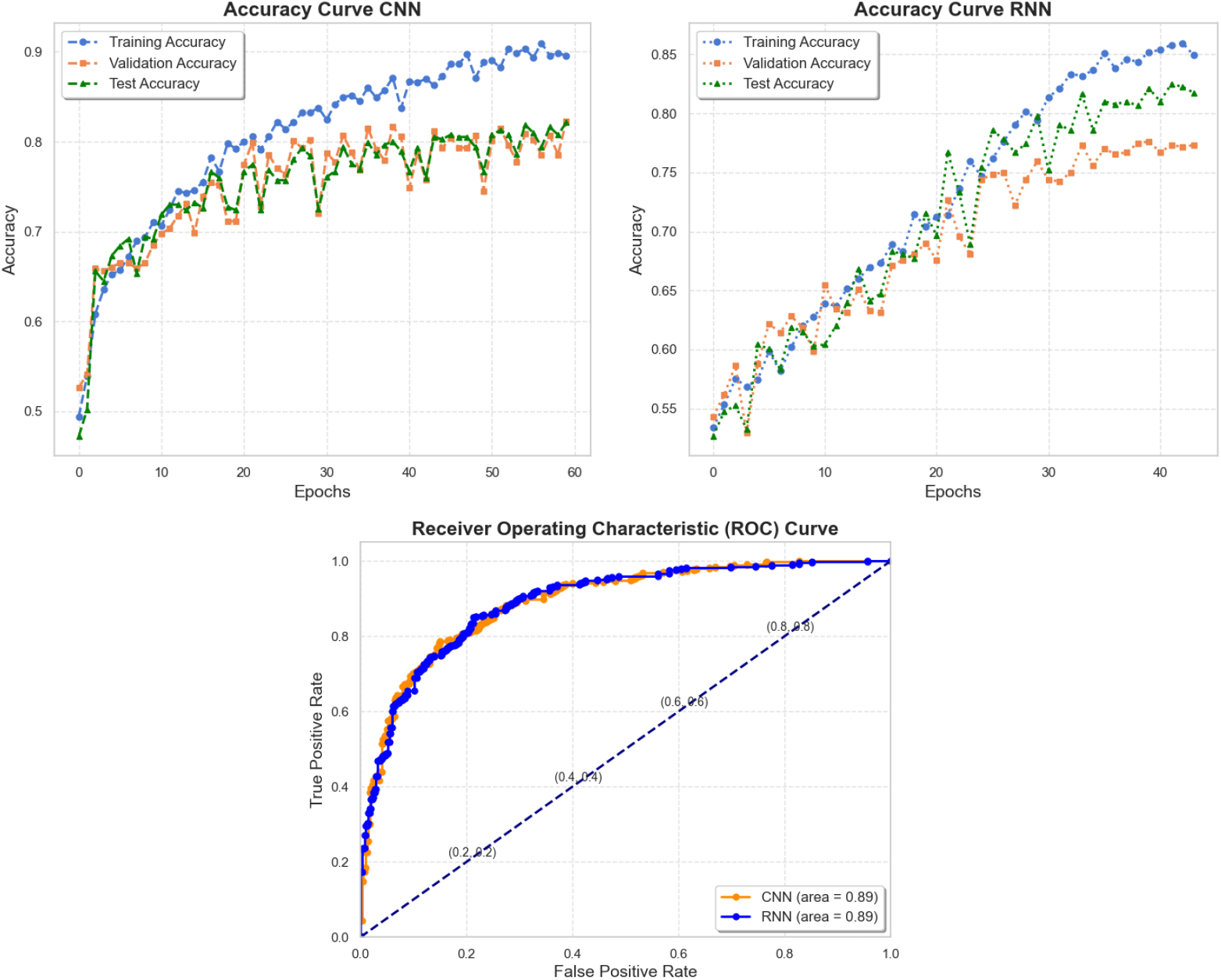
The training, validation, and test accuracy curves, along with the ROC curve for both CNN and RNN models, demonstrate the performance of the two models.

## Conclusion

We developed a cell-based modeling approach to study CTC dynamics within the HUMD. This innovative device, which exhibits local heterogeneity but global homogeneity in its structure, allowed for the identification and classification of CTCs based on their unique physiological characteristics, such as size, elasticity, and shape, result in distinct trajectories within the device. Our research demonstrated that these characteristics result in distinct trajectories within the HUMD, which, when analyzed using machine learning methods, can be used as indicators of CTC phenotypes. This suggests that label-free techniques could be developed to distinguish between various CTC subtypes. The findings of this study validate our concept of designing an innovative microfluidic device that can provide invaluable insights into CTC characteristics and phenotypes, potentially transforming approaches to early cancer diagnosis and treatment.

Despite the promising results, several questions remain unanswered. For instance, how do the observed trajectories differ across a wider range of CTC subtypes, and how can these differences be systematically quantified? Furthermore, the role of other biophysical factors, such as surface protein expression and cellular metabolism, in influencing CTC trajectories within the HUMD requires further exploration.

Understanding the interplay between these factors and their impact on CTC behavior could lead to even more precise classification methods. Another open question concerns the translation of this technology from the lab experiment to clinical settings. How can the HUMD be adapted or scaled for routine diagnostic use? What are the challenges and obstacles in standardizing the device and the associated machine learning algorithms across different patient populations? Addressing these questions is crucial for realizing the full potential of this technology in clinical practice. These areas will be the focus of our future research.

## Models

The utilization of a cell-based model is fundamental when it comes to investigating blood flow in microfluidic devices [18–20]. Unlike simple homogeneous fluids, blood is a complex medium composed of diverse cell types, including red blood cells (RBCs), white blood cells, and platelets, all suspended in a fluid called plasma. These various components contribute to the unique rheological properties of blood, making it a non-Newtonian fluid with distinct characteristics under different flow conditions [28, 29, 33–35]. The complexity is particularly pronounced within our proposed HUMD [13, 18, 36], as it encompasses not only the interactions between the CTCs and the HUMD but also their interactions with the other blood cells and plasma. Moreover, another significant challenge in studying the microfluidic flow in HUMD is the phenomenon known as shear-thinning behavior, as illustrated in Figure 13. This behavior refers to the reduction in viscosity that occurs when a fluid is subjected to increasing shear rates within microchannels [31]. In the context of microfluidic flow, this shear-thinning effect can profoundly impact the flow dynamics, particularly in the confined spaces of microfluidic channels. Therefore, by incorporating a cell-based model that accounts for the diverse components of blood and considers shear-thinning behavior, we can better understand and predict the complex behavior of blood flow and CTC motions in microfluidic devices.

## Fluid model

Plasma is considered as an incompressible, Newtonian fluid [33, 35–39], and the Lattice Boltzmann method (LBM) is employed to solve the Navier-Stokes equations (NSE). Instead of tackling the NSE directly, the LBM employs discrete particle populations that move along a lattice in discrete time steps Δ*t*. The movement of these particle populations is governed by a discretized Lattice Boltzmann equation,

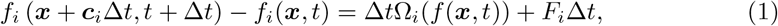

with the total number of populations given by *i* = 0, …, *q −* 1. *F*_*i*_ denotes the external forcing term. The particle populations *f*_*i*_ provide information about the concentration of particles with a specific velocity ***c***_*i*_ at a particular position ***x*** and time *t*. In each discrete time step Δ*t*, the populations move to adjacent lattice point *i* along a set of discrete velocity vectors ***c***_*i*_ with a lattice spacing of Δ*x*. This process, known as the streaming step, is depicted on the left-hand side of the LB equation. Additionally, the populations can interact with one another at the neighboring node, determined by the collision operator Ω_*i*_(*f* (***x***, *t*)) on the right-hand side of the LB equation. The operator redistributes the particles among the various populations *f*_*i*_ at each lattice site. The most straightforward collision operator utilized for the NS equations is derived from the BhatnagarGross-Krook (BGK) approximation and is expressed as [40] :

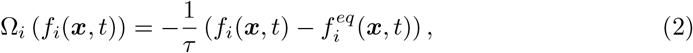

where *τ* is the relaxation time of the lattice Boltzmann fluid and 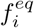 is the equilibrium population. The equilibrium populations 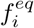 is given by

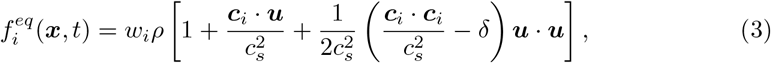

with weights *w*_*i*_ that belong to the chosen velocity set [34, 39, 41]. The ***u*** is the fluid macroscopic velocity. The collision operator relaxes the populations towards an equilibrium state 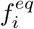, with a rate defined by the relaxation time *τ*. The relaxation time depends on the normalized speed of sound 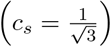 and the LBM kinematic viscosity *ν*_*lbm*_ by 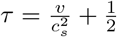. The parameter *τ* correlates with fluid viscosity. It establishes a connection between Lattice Boltzmann viscosity *ν*_*lbm*_, physical viscosity *ν*_*p*_, grid size *dx*, and time step *dt*.

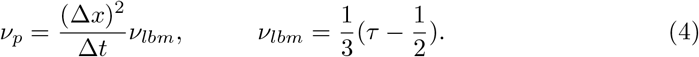

*ν*_*lbm*_ has a positive value for *τ >* 0.5. Now, combining them together we get:

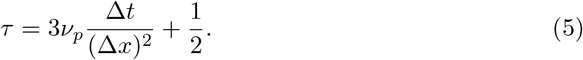

The driving force is incorporated via the term *F*_*i*_ :

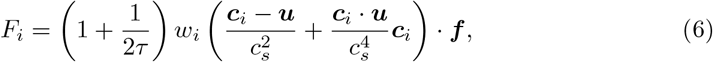

where ***f*** is the body force density [41]. This term is needed for the fluid-structure coupling of the fluid nodes to the particle membranes. The macroscopic NS equations can be derived which is achieved using the Chapman Enskog analysis. The zeroth and first velocity moment of the probability density function *f*_*i*_ give the macroscopic variables density *ρ* and momentum *ρu* of the fluid, respectively:

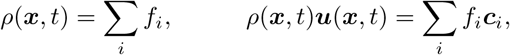

where the fluid velocity ***u*** is obtained as a weighted average of the particle velocity *c*_*i*_. The LBM offers the benefit of straightforward parallelization owing to its local dynamics. It consists of a linear streaming component (convection) and a non-linear collision operator that can be resolved locally.

## Constitutive model for cells

The 2*D* membrane model described in [19] is used to simulate the cells. This model incorporates various forces that influence the motion of the cellular membrane.

The link force operates along links between surface points and mimics the reaction to stretching and compression of the spectrin network, namely

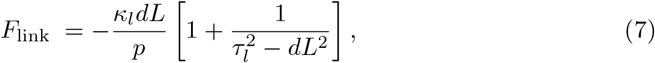

Where 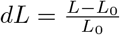 is the normal strain defined as the relative deviation from the equilibrium length *L*_0_, and *τ*_*l*_ specifies the maximum expandable length over the persistence length *p*.

The bending force acts between two adjacent surface elements, representing the membrane’s reaction to bending. This force is oriented along the normal direction of each surface, namely

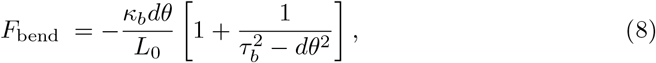

where *dθ* = *θ*_*i*_ *− θ*_0_. *τ*_*b*_ represents the limiting angle, which scales with the discretization length of the surface elements *L*_0_. The smallest representable curvature is fixed to 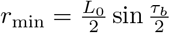.

The local surface conservation force operates within the vicinity of surface elements, capturing the combined surface response of the supporting spectrin network and the lipid bilayer of the membrane to stretching and compression. This force is exerted on all the vertices of each surface element, and it points towards the centroid of the corresponding surface element, namely

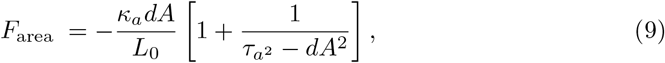

Where 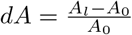.

The volume conservation force stands as the sole global factor and is employed to sustain the quasi-compressibility of the cell. It is applied at every surface vertex and it points towards the centroid of the cell, namely

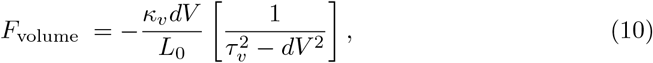

Where 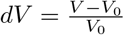, *τ*_*v*_ = 0.01 and *κ*_*v*_ = 20*k*_*B*_*T* is chosen to be a large but numerically still stable constant. The membrane constitutive model involves three independent parameters for cell representation: *κ*_*l*_, *κ*_*b*_, and *κ*_*a*_. These parameters are tuned to align with the mechanical characteristics in single cell experiments.

Other forces, such as the adhesive force due to the antibody-antigen reaction can be modeled based on the Lennard-Jones potential. Specifically, the attractive force is applied between the node representing the cell surface and the nodes on the surface of the micro-posts located within the microfluidic channel.

### Fluid structure interaction

The fluid problem in the Eulerian mesh is coupled with the cell membrane problem in the Lagrangian domain using the Immersed Boundary Method (IBM). The two-way coupling works in the following manner: the cell membrane exerts a force ***F*** (***X***, *t*) on the fluid, while the cell membrane feels the velocity of the fluid ***u***(***x***, *t*). Specifically, the force on the fluid ***f*** (***x***, *t*) at fluid node ***x*** and time *t* is obtained by spreading force ***F*** (***X***, *t*) using the following formula:

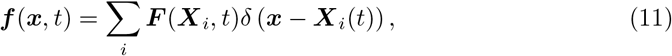

where ***X***_*i*_ denotes the coordinates of membrane node *i* and *δ* (***x*** *−* ***X***_*i*_(*t*)) is the Dirac delta function with a finite domain. The velocity ***U*** (***X***_*i*_, *t* + Δ*t*) on the membrane node ***X***_*i*_ is obtained by an interpolation step over the local fluid velocity field, using:

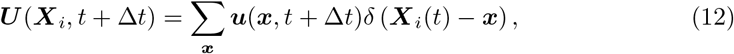

where ***u***(***x***, *t* + Δ*t*) is the velocity at fluid lattice coordinate ***x***. Membrane point ***X***_*i*_ moves due to advection can be calculated via the forward Euler scheme:

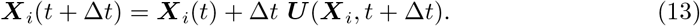

## Supporting information

**S1 Code Trajectory data and code** The cell trajectory data and machine learning code can be downloaded from https://github.com/imsanjoykb/vMDpcDI-CTC_Modeling.git

**S2 Code Cell based simulation code** The implementation and settings of our cancer cell simulation based on the Hemocell library [19] can be found here. https://github.com/qcutexu/CellBasedModeling.git

**S3 Video Case (A)** A movie showing the RBC tank treading motion under the shear rate 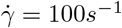.

**S4 Video. Case (B)** A movie showing the RBC tank treading motion under the shear rate 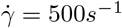.

**S5 Video. Case (B)** A movie showing a mixed tank treading and tumbling motions under the shear rate 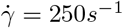.

**S6 Table.**
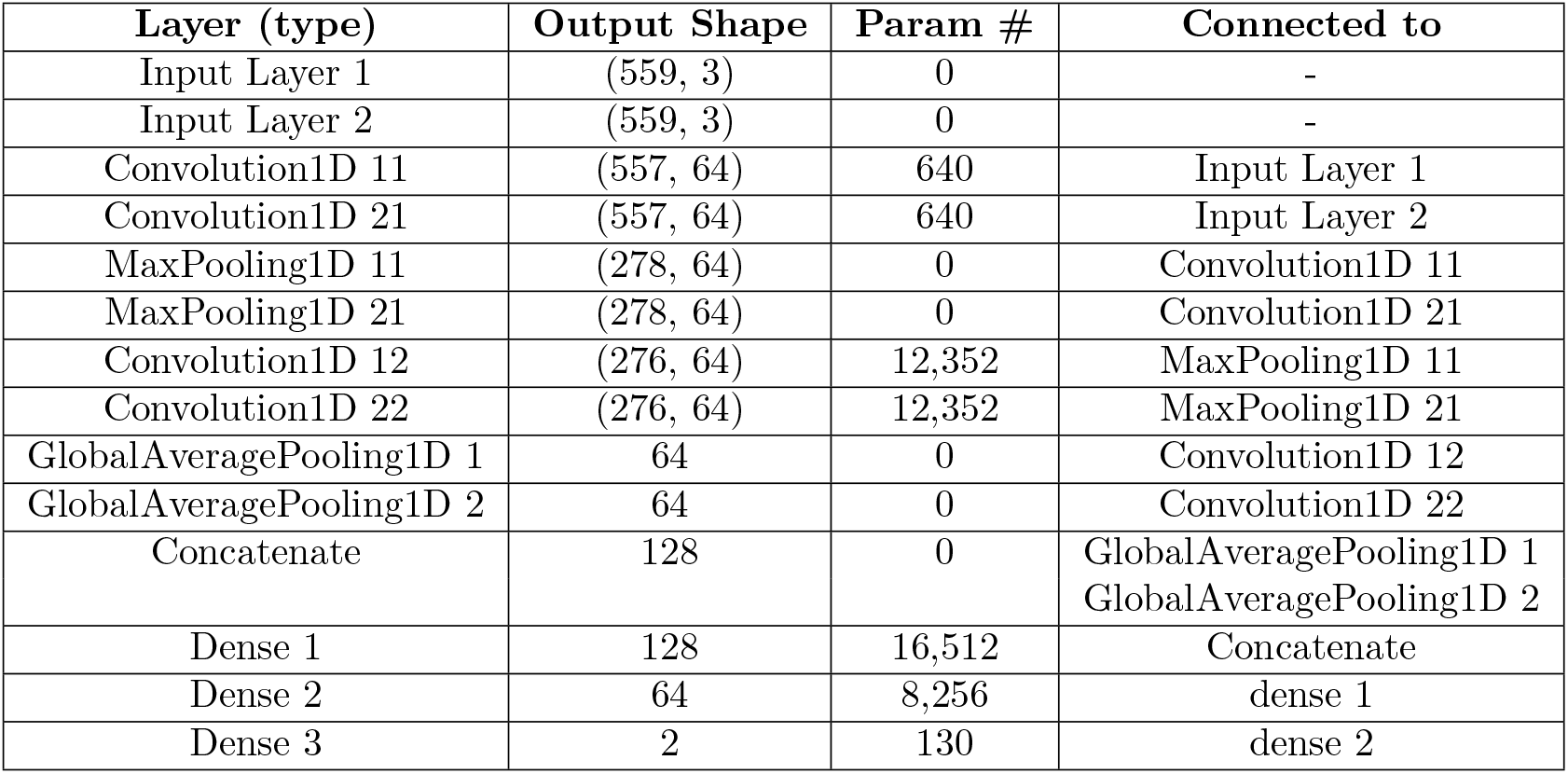
The configuration of RNN. The details of the recurrent layers and hidden layers are provided in the table below.

**S7 Table.** The configuration of CNN. The details of the convolutional layers and hidden layers are provided in the table below.

| Layer (type) | Output Shape | Param # |
| --- | --- | --- |
| Bidirectional 1 | (None, 559,256 ) | 104,448 |
| Batch normalization 1 | (None, 559, 256) | 1,024 |
| Dropout 1 | (None, 559,256 ) | 0 |
| Bidirectional 2 | (None, 559,128 ) | 123,648 |
| Batch normalization 2 | (None, 559,128 ) | 512 |
| Dropout 2 | (None, 559,128 ) | 0 |
| Bidirectional 3 | (None, 64) | 31,104 |
| Batch normalization 3 | (None, 64) | 256 |
| Dropout 3 | (None, 64) | 0 |
| Dense 1 | (None, 16) | 1,040 |
| Dense 2 | (None, 2) | 34 |

## Acknowledgments

This research was partially funded by the National Science Foundation through grants DMS-2247000 (Canic), DMS-2247001 (Wang), and CBET-1935792 (Li), as well as by the National Institutes of Health (IMAT, Grant No. 1R21CA240185-01). The authors also acknowledge the High-Performance Computing Center (HPCC) at Texas Tech University for providing computational resources that have contributed to the research results reported within this paper (URL: http://www.hpcc.ttu.edu). The authors would like to express their gratitude to Karl Gardner at Texas Tech University for sharing the experimental data used for our validation purposes.

